# Regulation of blood cell transdifferentiation by oxygen sensing neurons

**DOI:** 10.1101/2020.04.22.056622

**Authors:** Sean Corcoran, Anjeli Mase, Yousuf Hashmi, Debra Ouyang, Jordan Augsburger, Thea Jacobs, Katelyn Kukar, Katja Brückner

**Affiliations:** Eli and Edythe Broad Center of Regeneration Medicine and Stem Cell Research; Department of Cell and Tissue Biology; Cardiovascular Research Institute, University of California San Francisco, San Francisco, CA

**Keywords:** *Drosophila melanogaster*, transdifferentiation, hematopoiesis, microenvironment, hemocyte, macrophage, plasmatocyte, crystal cell, sensory neurons, sensory cones, atypical guanylyl cyclase, oxygen sensing, hypoxia

## Abstract

Transdifferentiation generates specialized cell types independent of stem or progenitor cells. Despite the unique process, it remains poorly understood how transdifferentiation is regulated in vivo. Here we reveal a mechanism of environmental control of blood cell transdifferentiation in a *Drosophila* model of hematopoiesis. Functional lineage tracing provides evidence for transdifferentiation from macrophage-like plasmatocytes to crystal cells that execute melanization. Interestingly, this transdifferentiation is promoted by neuronal activity of a specific subset of sensory neurons, in the caudal sensory cones of the larva. Crystal cells develop from plasmatocyte clusters surrounding the sensory cones, triggered by environmental conditions: oxygen sensing, and the atypical guanylyl cyclase Gyc88E specifically expressed in the sensory cone neurons, drive plasmatocyte-to-crystal cell transdifferentiation. Our findings reveal an unexpected functional and molecular link of environment-monitoring sensory neurons that govern blood cell transdifferentiation in vivo, suggesting similar principles in vertebrate systems where environmental sensors and blood cell populations coincide.

**Highlights:**

- Functional lineage tracing reveals in vivo transdifferentiation in a *Drosophila* model of hematopoiesis
- Active sensory neurons of the caudal sensory cones promote blood cell transdifferentiation in the *Drosophila* larva
- Environmental oxygen sensing and atypical guanylyl cyclase activity in sensory cone neurons drive blood cell transdifferentiation

## Introduction

### A Drosophila model of blood cell transdifferentiation

The phenomenon of transdifferentiation has been noted in a variety of species (Cieslar-Pobuda et al., 2017; Graf, 2011; Reid and Tursun, 2018), yet its in vivo regulation remains poorly understood. In the vertebrate blood cell system, reports of transdifferentiation have for the most part been limited to cell culture systems and experimental manipulations. For example, C/EBP (CCAAT/enhancer-binding protein) transcription factors drive transdifferentiation of vertebrate B cells to macrophages (Di Tullio et al., 2011; Xie et al., 2004), B lymphoma and leukemia cell lines to macrophages (Rapino et al., 2013), and B cells to Granulocyte-Macrophage Precursors (Cirovic et al., 2017). Similarly, manipulation of key transcription factors such as FLI1 and ERG results in transdifferentiation of erythroblasts to megakaryocytes (Siripin et al., 2015), and deletion of the BAF Chromatin Remodeling Complex Subunit Bcl11b triggers T cell transition to NK cells (Li et al., 2010). Transdifferentiation of lymphoid and myeloid cells has been modeled mathematically (Collombet et al., 2017). However, in vivo, the underlying cellular and molecular mechanisms of blood cell transdifferentiation during development and homeostasis and the role of the environment remain elusive.

To investigate principles of in vivo transdifferentiation in the hematopoietic system, we turned to a model in *Drosophila melanogaster. Drosophila* offers proven parallels to the two major lineages that produce myeloid blood cells in vertebrates (Davies et al., 2013; Perdiguero and Geissmann, 2016; Sieweke and Allen, 2013), with its two myeloid lineages of blood cells, or hemocytes (Gold and Brückner, 2014, 2015; Holz et al., 2003), (1) the embryonic lineage of hemocytes that proliferate as differentiated cells in hematopoietic pockets of the larval body wall, and resemble vertebrate tissue macrophages, and (2) the progenitor-based lymph gland lineage (Banerjee et al., 2019). Both *Drosophila* blood cell lineages produce at least three differentiated blood cell types: macrophage-like plasmatocytes, crystal cells that mediate melanization, and lamellocytes, large immune cells specialized for encapsulation (Banerjee et al., 2019; Gold and Brückner, 2014, 2015). During larval stages, embryonic-lineage plasmatocytes show signs of fate changes to other blood cell types: Transdifferentiation to lamellocytes occurs in response to immune challenges (Markus et al., 2009), and transdifferentiation to crystal cells was suggested even under steady state conditions (Leitao and Sucena, 2015). However, it remains unclear what are the anatomical requirements and environmental inputs that regulate plasmatocyte-to-crystal cell transdifferentiation in vivo. Hematopoietic sites of the *Drosophila* larva contain sensory neuron clusters of the peripheral nervous system (PNS) that serve as microenvironments for plasmatocyte survival, proliferation and localization (Gold and Brückner, 2014, 2015; Makhijani et al., 2017; Makhijani et al., 2011; Makhijani and Brückner, 2012). Considering this, we investigated the role of these specialized hematopoietic pockets in hemocyte transdifferentiation.

## Results

### Phagocytic plasmatocytes transdifferentiate to crystal cells

First we examined the formation of crystal cells and their anatomical locations during larval development. We quantified crystal cells using a traditional way of labeling crystal cells based on their expression of prophenoloxidases (Corcoran and Brückner, 2020; Rizki and Rizki, 1959), enzymes responsible for melanization, the main immune function of crystal cells (Bidla et al., 2009; Dudzic et al., 2015; Lu et al., 2014). Induction of melanization blackens crystal cells (Corcoran and Brückner, 2020; Rizki and Rizki, 1959) and marks similar cell populations as the crystal cell reporter *lozenge-GAL4* (*lz-GAL4; UAS-GFP*) (SupplFig. 1 A, B). During larval development, crystal cell numbers increase slowly over the first and second larval instar stages, but expand more rapidly during the third instar stage (Fig. 1 A), largely mirroring the exponential increase of plasmatocytes during larval development (Fig 1 B) (Makhijani et al., 2011; Petraki et al., 2015). To visualize the locations of crystal cells and plasmatocytes, we coexpressed two fluorescent protein reporters (for crystal cells *BcF2-GFP* (Tokusumi et al., 2009), and for plasmatocytes *Hml*Δ*-DsRed* (Makhijani et al., 2011)). Interestingly, crystal cells are strongly enriched in a cluster in the terminal segment of the *Drosophila* larva, a region where plasmatocytes are also known to accumulate (Fig. 1 C-C’’, D-D’) (Makhijani et al., 2011). Crystal cells are occasionally also found in other hematopoietic pockets, and in dorsal vessel-associated clusters where floating hemocytes accumulate (Cevik et al., 2019; Petraki et al., 2015).

**Figure 1.**
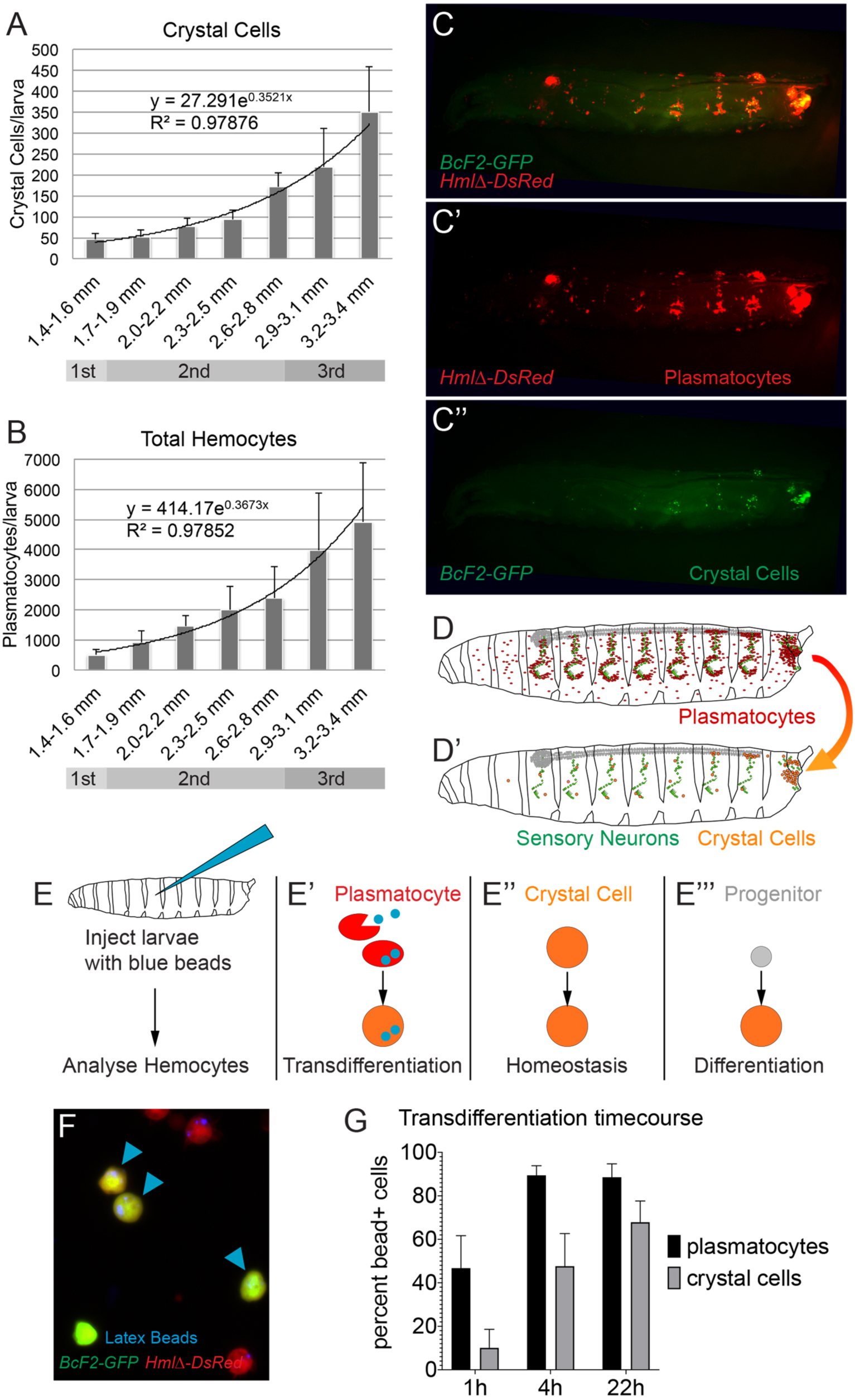
Crystal cells in the *Drosophila* larva are generated by transdifferentiation. (A) Development of crystal cell numbers over time. Crystal cells per larva, assessed by melanization, relative to larval size and developmental stage. Genotype is *w*^*1118*^, n=155. Mean and standard deviation, regression analysis. (B) Development of total hemocyte numbers over time. Total hemocytes per larva, genotype is *HmlΔGAL4, UAS-GFP; He-GAL4*; n=107. Mean and standard deviation, regression analysis. (C-C’’) Localization of plasmatocytes and crystal cells in the 3^rd^ instar larva,; genotype is *BcF2-GFP/ Hml*Δ*-DsRed*; plasmatocytes labeled by *Hml*Δ*-DsRed* (red); crystal cells labeled by *BcF2-GFP* (green); lateral view, posterior right. (D-D’) Model of distribution of plasmatocyte (red) and crystal cells (orange) in *Drosophila* larva, relevant sensory neuron clusters of the hematopoietic pockets (green); lateral view, posterior right. (E-E’’’) Scheme of phagocytosis lineage tracing assay. Blue fluorescent latex beads are injected into early 3^rd^ instar larvae, cells are released after indicated incubation times; fraction of cells containing phagocytosed beads are determined. Possible outcomes are depicted with crystal cells in orange, plasmatocytes in red, progenitors in grey. (F) Sample image of analyzed cells; genotype *BcF2-GFP/ Hml*Δ*-DsRed*; crystal cells in green, plasmatocytes in red; blue arrowheads indicate crystal cells with incorporated blue beads. (G) Quantification of samples as in (G); fraction of plasmatocytes and crystal cells containing blue beads at time 1h, 4, 22h after injection; n=21; mean and standard deviation.

Since previous studies suggested plasmatocyte-to-crystal cell transdifferentiation solely based on live imaging (Leitao and Sucena, 2015), we sought an independent functional lineage tracing approach. Asking whether crystal cells derive from undifferentiated progenitors or differentiated, phagocytically active plasmatocytes, we traced cells based on the unique ability of differentiated plasmatocytes to phagocytically uptake fluorescently labeled beads (Fig. 1 E-E’’’). We injected *Drosophila* larvae expressing fluorescent reporters for plasmatocytes and crystal cells with blue fluorescent latex beads. Injected larvae were incubated in a time course, followed by the release of hemocytes and quantification of the relative fractions of phagocytosis-labeled plasmatocytes and - crystal cells (Fig. 1 F, G). The fraction of blue bead positive plasmatocytes quickly reached saturation (∼50% at 1h, ∼90% at 4h and 22h). In contrast, crystal cells were labeled by blue beads with a significant time delay (<10% at 1h, ∼ 50% at 4h, ∼70% at 22h) (Fig. 1 G). Together with previous reports that suggested crystal cells are not capable of phagocytosis or proliferation (Lanot et al., 2001; Leitao and Sucena, 2015; Tattikota et al., 2019) this supports a model of plasmatocyte-to-crystal cell transdifferentiation, in which crystal cells derive from phagocytically active plasmatocyes, rather than from undifferentiated, phagocytosis-incompetent progenitors.

### Sensory neuron activity promotes crystal cell transdifferentiation

Next, we investigated the anatomical locations of crystal cells and plasmatocytes relative to sensory neurons, using fluorescent reporters and live imaging, including the sensory neuron specific driver *21-7-GAL4, UAS-CD8-GFP* (Song et al., 2007) (Fig. 2 A). We found that both plasmatocytes and crystal cells colocalize with sensory neurons, with one important difference: plasmatocytes are found in all hematopoietic pockets (Fig. 2 A), while crystal cells are mainly localized in the terminal hematopoietic pocket of the larva (Fig. 2 B).

**Figure 2.**
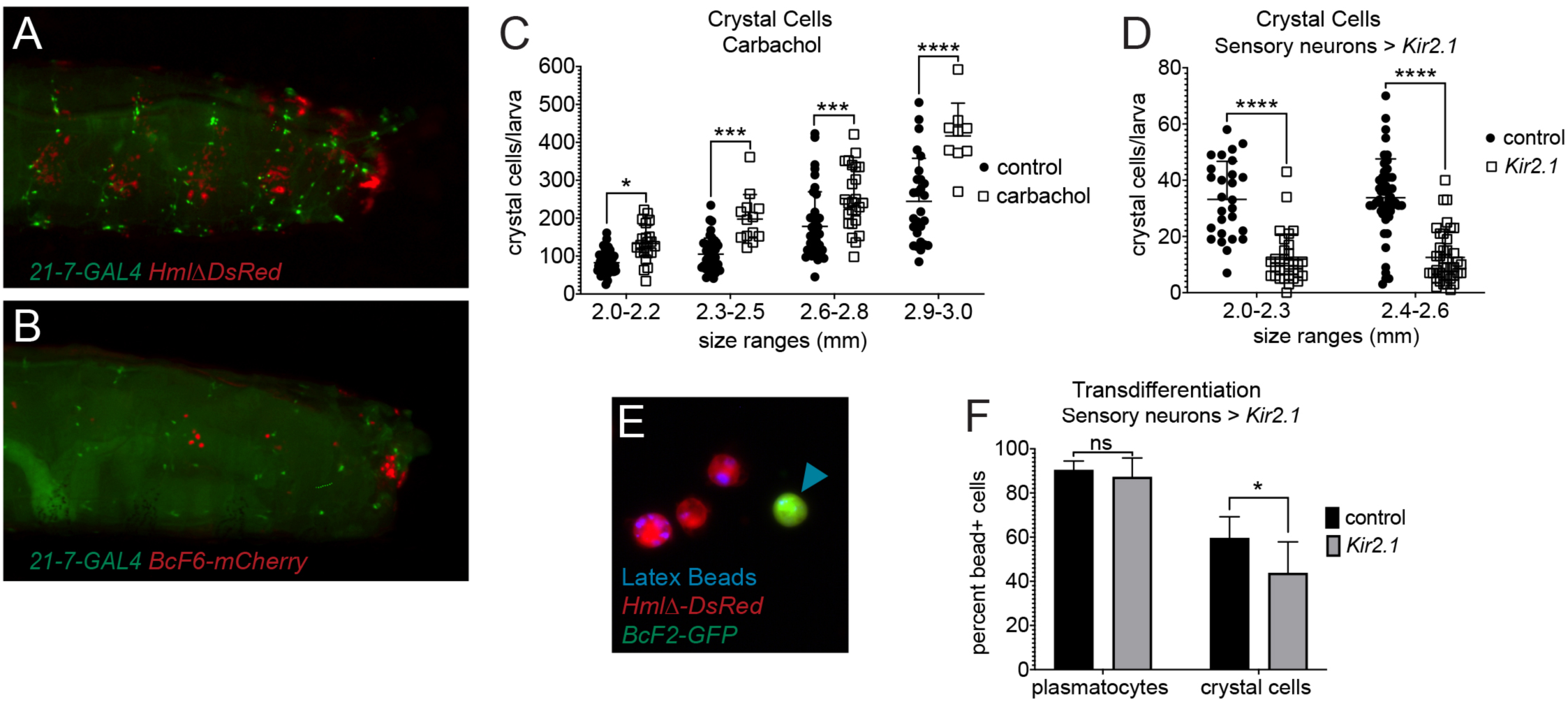
Sensory Neuron activity regulates transdifferentiation to crystal cells. (A) Plasmatocytes colocalize with sensory neurons in all hematopoietic pockets; genotype *21-7-GAL4, UAS-CD8-GFP, Hml*Δ*DsRed/CyO* ; lateral view, posterior right. (B) Crystal cells also colocalize with sensory neurons but are mainly found in a cluster at the caudal end of the larva; genotype *21-7-GAL4, UAS-CD8-GFP/*+; *BCF6-mCherry/*+; lateral view, posterior right. (C) Treatment of larvae with the AchR agonist carbachol to mimic sensory neuron activation, treatment for 4h; genotype is *yw*; quantification of crystal cells by melanization; control n=65 (+45); carbachol n=34 (+50). Individual value plot with mean and standard deviation, two-way ANOVA. (D) Transient silencing of sensory neurons, quantification of crystal cells by melanization; genotypes are experiment *21-7-GAL4, UAS-CD8GFP, Hml*Δ*-DsRed/ UAS-Kir2.1; tubGAL80ts/* + and control *21-7-GAL4, UAS-CD8GFP, Hml*Δ*-DsRed/* +; *tubGAL80ts/* +. Larvae induced at 29°C for 22h. Mean and standard deviation, two-way ANOVA. (E, F) Phagocytosis lineage tracing, genotypes are experiment *21-7-GAL4, UAS-CD8GFP, Hml*Δ*-DsRed/ UAS-Kir2.1; BcF6-GFP/ tubGAL80ts*, and control *21-7-GAL4, UAS-CD8GFP, Hml*Δ*-DsRed/* +; *BcF6-GFP/ tubGAL80ts*. (E) Sample image of analyzed cells, plasmatocytes (red), crystal cells (green), injected beads (blue). (F) Quantification of samples as in (E); fraction of plasmatocytes and crystal cells containing blue beads, experiment n=12 and control n=12. Mean and standard deviation, two-way ANOVA.

Given this colocalization with sensory neurons, we asked whether sensory neuron activity has an effect on crystal cell generation. To mimic activation of sensory neurons, we exposed larvae to the acetylcholine receptor agonist carbamoycholine (carbachol). Carbachol exposure over 4 hours resulted in a moderate increase of crystal cell numbers (Fig. 2 C). More specifically and complementary to this, we silenced sensory neurons by transient expression of *Kir2.1*, an inward rectifying K+ channel that causes neuron hyperpolarization (Baines et al., 2001). Interestingly, *Kir2.1* expression over 22 hours caused a dramatic drop in crystal cell numbers (Fig. 2 D). There was no significant effect of neuronal silencing on total hemocyte numbers under comparable conditions (SupplFig. 2 and (Makhijani et al., 2017)). To address whether sensory neuron silencing by Kir2.1 affects plasmatocyte to crystal cell transdifferentiation, we performed phagocytosis lineage tracing. Indeed, transient neuronal silencing affects the fraction of crystal cells derived from blue bead labeled plasmatocytes, while the ability of plasmatocytes to phagocytose remains the same (Fig. 2 E, F). Taken together, our findings suggest that sensory neuron activity promotes transdifferentiation of plasmatocytes to crystal cells.

### Crystal cells colocalize with, and require, sensory neurons of the sensory cones

To gain more insight, we focused on the caudal cluster of crystal cells. Closer inspection of combined fluorescent reporters for crystal cells and sensory neurons showed that crystal cells colocalize particularly well with sensory organs of the sensory cones (Fig. 3 A-A’’ and C-D’’), protruding structures that are grouped around the posterior spiracles, the terminal tubes of the tracheal system (Hayashi and Kondo, 2018; Kuhn et al., 1992). Plasmatocytes accumulate in a large cluster around the sensory cones already in the 2^nd^ instar larva (Fig. 3 B-B’’), which seems to foreshadow the pattern and abundance of crystal cells in the 3^rd^ instar larva (Fig. 3 D-D’’). We therefore hypothesized that many plasmatocytes of the 2^nd^ instar larva may transdifferentiate to crystal cells, as is apparent in the 3^rd^ instar.

**Figure 3.**
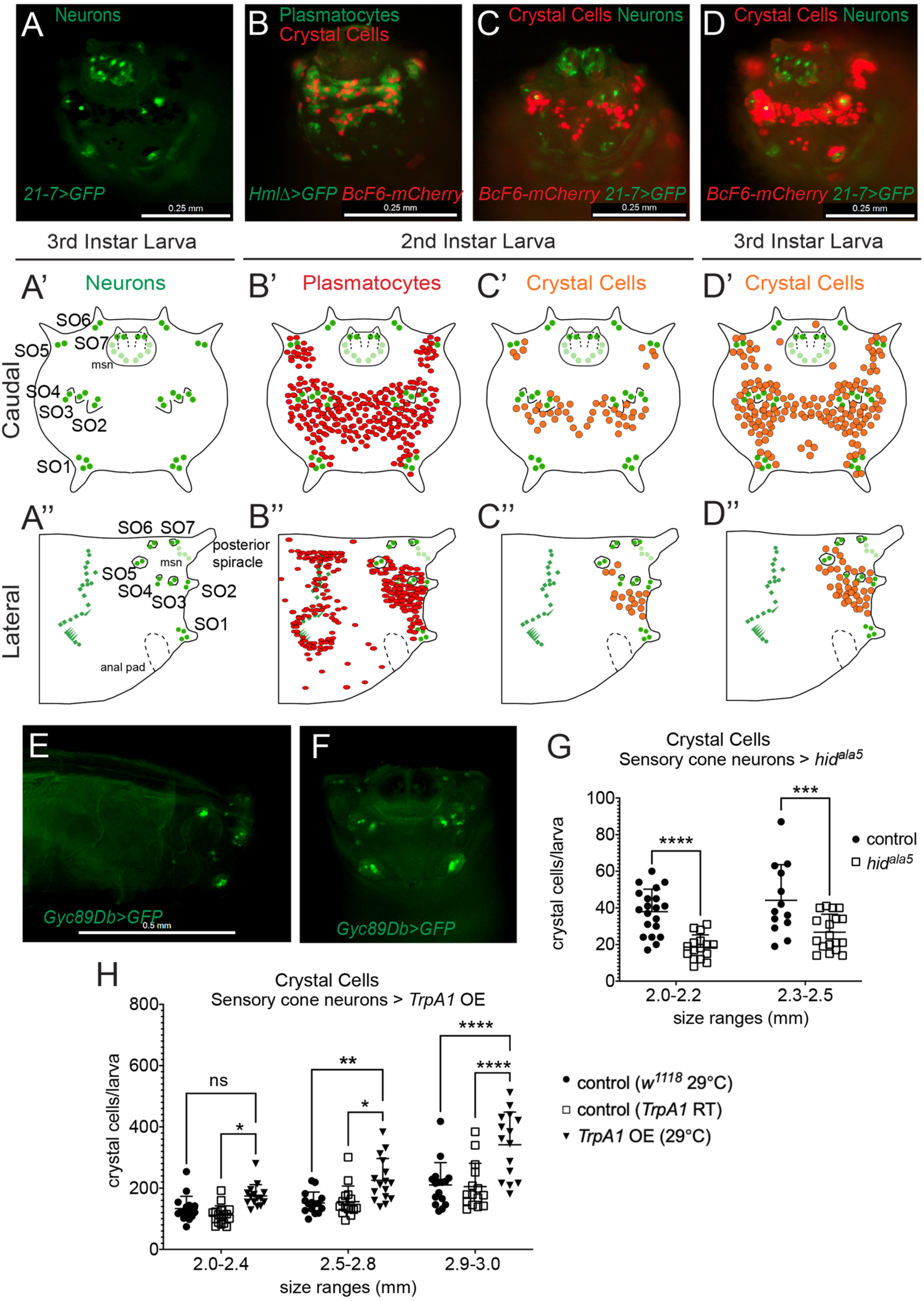
Crystal cells are clustered around the sensory cones and are promoted by sensory cone neurons. (A-D) Localization of sensory neurons, plasmatocytes and crystal cells, caudal view of larvae; scale bars 0.25mm. (A) Sensory neurons (green), genotype *21-7-GAL4, UAS-CD8-GFP, Hml*Δ*-DsRed/CyO*; 3rd instar larva. (B) Plasmatocytes (green) and crystal cells (red), genotype *Hml*Δ*-GAL4, UAS-GFP; BcF6-mCherry*; 2^nd^ instar larva. (C) Crystal cells (red) and sensory neurons (green), genotype *21-7-GAL4, UAS-CD8-GFP/*+; *BcF6-mCherry/*+; 2^nd^ instar larva. (D) Crystal cells (red) and sensory neurons (green), *21-7-GAL4, UAS-CD8-GFP/*+; *BcF6-mCherry/*+; 3rd instar larva. (A’-D’) Models corresponding to (A-D), respectively, plasmatocytes red, crystal cells orange, sensory neurons green; caudal view. (A’’-D’’) Models, lateral view, corresponding to (A-C and A’-C’), respectively; lateral view. (E) *Gyc89Db-GAL4* driver expressing GFP in sensory cone neurons (green), lateral view, genotype is *UAS-GFP/*+; *Gyc89Db-GAL4/*+; lateral view, posterior right; scale bar 0.5mm. (F) Larva as in (E), caudal view. (G) Ablation of sensory cone neurons affects crystal cells; quantification of crystal cells per larva by melanization. Genotypes are experiment *UAS-Hid ala5/*+; *Gyc89Db-GAL4/tubGAL80ts*, n=34 and control *Gyc89Db-GAL4/tubGAL80ts*, n=34. Crosses were raised at 18°C and temperature shifted to 29 °C for 16 h. Individual value plot with mean and standard deviation, two-way ANOVA. (H) Transient activation of TrpA1 in sensory cone neurons; quantification of crystal cells per larva by melanization. Genotypes are experiment *UAS-TrpA1/* +; *Gyc89Db-GAL4, /* +, n=46, and control *Gyc89Db-GAL4/*+, n=48; in addition, one experiment F1 cohort *UAS-TrpA1/* +; *Gyc89Db-GAL4, /* +, n=47, was maintained as uninduced control at RT. Crosses were raised at RT and temperature shifted to 29°C for 4 hours;. Individual value plot with mean and standard deviation, two-way ANOVA.

Given the intriguing colocalization of crystal cells and sensory neurons and the dependence of blood cell transdifferentiation on neuronal activity, we investigated the specific requirement of sensory cone neurons for crystal cell production. A group of genes specifically expressed in sensory cone neurons are atypical guanylyl cyclases, *Gyc88E, Gyc89Da*, and *Gyc89Db* (Vermehren-Schmaedick et al., 2010). Using the driver *Gyc89Db-GAL4* (Vermehren-Schmaedick et al., 2010) (Fig. 3 E, F), we ablated sensory cone neurons by transiently expressing the proapoptotic gene *head involution defect (Hid)* in its non-repressible version (*Hid* ^*ala5*^) (Bergmann et al., 2002). Ablation of sensory cone neurons did not affect larval viability, but it strongly reduced crystal cell numbers (Fig. 3 G). Conversely, ectopic activation of sensory cone neurons by specific expression and transient induction of the heat-induced cation channel TrpA1 (Hamada et al., 2008) caused a significant increase in crystal cells (Fig. 3 H). Taken together, we conclude that crystal cells and their precursor plasmatocytes are strongly enriched at the sensory cones of the larva. Sensory cone neurons, and their activity, are required for crystal cells.

### Oxygen sensing through atypical guanylyl cyclases drives plasmatocyte-to-crystal cell transdifferentiation

The atypical guanylyl cyclase Gyc88E is a cytoplasmic oxygen sensor that acts as homodimer or forms heterodimers with related guanylyl cyclases such as Gyc89Da or Gyc89Db, (Huang et al., 2007; Morton, 2004; Morton et al., 2005; Vermehren et al., 2006). Gyc88E complexes, which are active already at ambient oxygen levels, generate the second messenger cyclic GMP (cGMP) that in turn activates neurons (Morton, 2004; Morton et al., 2008). Since we found that sensory cone neurons expressing these Gycs are required for crystal cell formation, we decided to test Gyc function itself in relation to crystal cell transdifferentiation. When the obligatory subunit, Gyc88E, was silenced in the sensory cone neurons, crystal cell numbers were reduced (Fig. 4 A). Phagocytosis lineage tracing confirmed that Gyc88E function is required for transdifferentiation of plasmatocytes to crystal cells, as the fraction of crystal cells carrying blue beads was significantly reduced in *Gyc88E* RNAi knockdowns, while phagocytosis by plasmatocytes remained the same (Fig. 4 B). Given the role of Gycs as oxygen sensors, we next investigated the effect of varying atmospheric oxygen concentrations on crystal cell formation. Assessing crystal cell numbers per larva as readout, we exposed larvae to reduced levels of oxygen (8%, or 5%) for 6 hours. Compared to normoxia (∼21% oxygen), 8% hypoxia caused mild reduction in crystal cells and 5% oxygen caused a significant drop in crystal cells, while total hemocyte numbers remained unaffected (SupplFig. 3 A, B, Fig. 4 C). Following up on the robust results obtained under 5% oxygen, we determined whether hypoxia affects blood cell transdifferentiation, performing phagocytosis lineage tracing for 6 hours under hypoxic conditions (5% O_2_) and normoxia. Interestingly, hypoxia phenocopies silencing of Gyc88E function in sensory cone neurons, resulting in a significant reduction of blue beads in crystal cells, while phagocytosis levels of plasmatocytes remain unaffected (Fig. 4 D). Our data demonstrate a role for oxygen and the atypical guanylyl cyclase Gyc88E, likely though its link to activation of the sensory cone neurons, in the promotion of plasmatocyte-to-crystal cell transdifferentiation (Fig. 4 E). This model supports the kinetics of crystal cells following plasmatocyte expansion, based on the transition of plasmatocytes to crystal cells in particular in the proximity of sensory cone neurons (SupplFig. 4 A).

**Figure 4.**
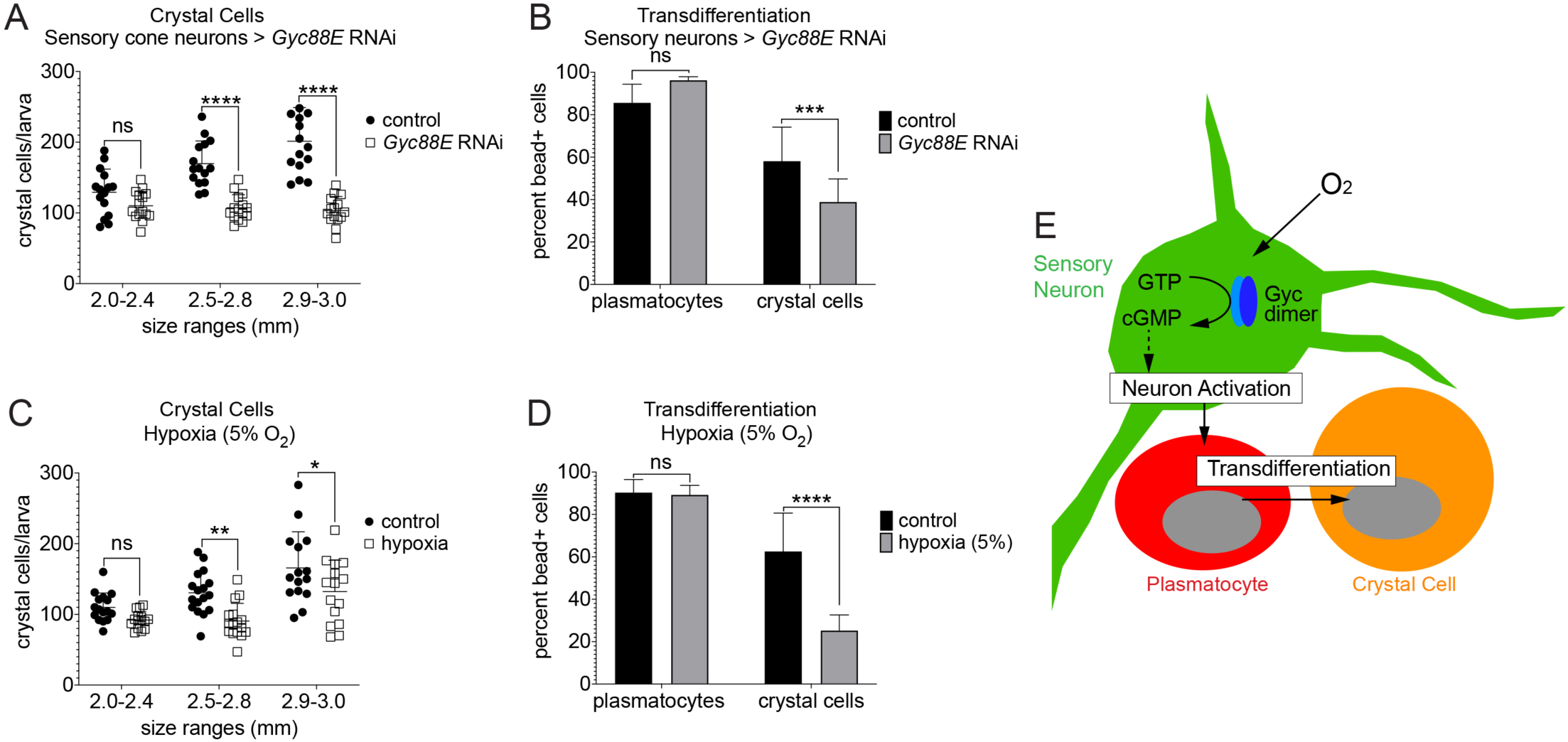
Oxygen sensing through Gycs in sensory cone neurons drives plasmatocyte-to-crystal cell transdifferentiation. (A) RNAi silencing of *Gyc88E* in sensory cone neurons results in reduced crystal cell numbers determined by melanization; genotyes are experiment *Gyc89Db-GAL4/UAS-Gyc88ERNAi*, n=45 and control *Gyc89Db-GAL4/*+. n=45. Individual value plot with mean and standard deviation, two-way ANOVA. (B) Phagocytosis lineage tracing, effect of *Gyc88E* RNAi in sensory cone neurons on transdifferentiation; genotypes are experiment *21-7-GAL4, UA5-CD8-GFP, Hml*Δ*-DsRed/*+; *BcF6-GFP/ UAS-Gyc88ERNAi*, n=11 and control *21-7-GAL4, UA5-CD8-GFP, Hml*Δ*-DsRed/*+; *BcF6-GFP/*+, n=16. Bar chart with mean and standard deviation, two-way ANOVA. (C) Effect of hypoxia (5% O2) on crystal cell number per larva determined by melanization; genotype is *w*^*1118*^; hypoxia n=46 and normoxia n=48. Individual value plot with mean and standard deviation, two-way ANOVA. (D) Phagocytosis lineage tracing, effect of hypoxia (5% O2) on transdifferentiation; genotype is *Hml*Δ*-GAL4, UAS-GFP; BcF6-mCherry*; hypoxia n=14 and normoxia n=15. Bar chart with mean and standard deviation, two-way ANOVA. (E) Model. Sensory cone neurons require oxygen and cytoplasmic Gyc88E to drive crystal cell formation. Gyc88E converts GTP to cGMP, leading to downstream signaling and neuronal activation. Through a currently unknown mechanism, long-term hypoxia negatively affects this process. Active neurons induce plasmatocyte-to-crystal cell transdifferentiation.

## Discussion

Our work has identified an unexpected mechanistic link between oxygen sensing and blood cell transdifferentiation, which is facilitated through a particular set of sensory neurons and an intracellular atypical guanylyl cyclase. This new paradigm inspires the search for similar principles of neuronally controlled blood cell transdifferentiation that responds to environmental conditions in other species including humans.

Transdifferentiation results in the conversion of one differentiated cell type to another. In some systems, new differentiated cell types arise after de-differentiation to a transient pluripotent intermediate (Graf, 2011; Pesaresi et al., 2019; Reid and Tursun, 2018). Our work supports a model of direct transdifferentiation based on fluorescent reporters and phagocytosis lineage tracing.

Independent approaches also support a model of continuous blood cell transdifferentiation from plasmatocytes to crystal cells through progressive states, based on single cell RNAseq pseudotime lineage analysis (Tattikota et al., 2019). Transdifferentiation may be the fastest and most efficient way for animals to shape the composition of their blood cell pool according to environmental conditions such as oxygen levels and potentially other inputs.

Gyc intracellular oxygen sensors mediate oxygen detection in sensory neurons (Vermehren et al., 2006), similar to other oxygen sensing mechanisms known in vertebrate neurons and other sensory cells (Caravagna and Seaborn, 2016; Pokorski et al., 2016). In contrast, HIF (hypoxia inducible factor) transcription factors regulate target genes in response to low oxygen conditions in a variety of cell types (Gorr et al., 2006; Majmundar et al., 2010). Hypoxia, through HIF, regulates mammalian hematopoiesis, lymphopoiesis and erythropoiesis (Chabi et al., 2019; Haase, 2013; Imanirad and Dzierzak, 2013). The *Drosophila* HIF1a *sima* (*similar*) plays a role in crystal cell formation in the *Drosophila* lymph gland (Mukherjee et al., 2011), however this effect is independent of HIF1β and hypoxia target genes. Instead, formation of *Drosophila* crystal cells has been linked to Notch signaling at various developmental stages (Duvic et al., 2002; Krzemien et al., 2007; Lebestky et al., 2000; Lebestky et al., 2003; Leitao and Sucena, 2015; Mukherjee et al., 2011). It remains to be determined whether oxygen sensing neurons and/or the signal receiving plasmatocytes are connected to Notch signaling or an independent pathway that governs the fate switch of plasmatocytes to crystal cells.

Linking oxygen sensing to crystal cell formation via neuronal activity involves Gyc88E and likely additional factors that modulate its activity under long-term hypoxia. While hypoxia in the range of minutes enhances Gyc88E activity and induces short-term behavioral responses in *Drosophila* larvae, additional regulatory factors such as nitric oxide (NO) are known to dampen Gyc88E activity (Huang et al., 2007) and could be responsible for the reduction in crystal cell transdifferentiation we observe. Interestingly, NO is known to be induced by long-term hypoxia and shows particularly high levels in the sensory cones (Wingrove and O’Farrell, 1999). Additionally or alternatively, long-term hypoxia could affect Gyc88E activity through refractory mechanisms, interaction with additional regulatory interaction partners (Ding et al., 2020), links to alternative downstream effectors, or even alternative function. Under short-term hypoxia, Gycs generate cGMP which acts through the cyclic nucleotide gated channel A (CNGA) (Morton et al., 2008; Vermehren-Schmaedick et al., 2010). CNG channels mediate influx of calcium ions, among others, resulting in consecutive activation of calmodulin/CaMK signaling and sensory transduction (Kaupp and Seifert, 2002; Pifferi et al., 2006).

While it is difficult to determine the effects of short-term hypoxia on the comparably slow process of blood cell transdifferentiation, the key steps of signaling downstream of Gyc88E in blood cell transdifferentiation and the quantitative and/or qualitative differences of signaling under conditions of long-term hypoxia remain to be seen.

Regulation of blood cell transdifferentiation by sensory cone neurons offers advantages of immediate sensitivity and may further integrate the response to a variety of environmental conditions. Sensory cone neurons are in contact with the surrounding atmosphere, based on social burying during feeding in which larvae expose their caudal ends with the sensory cones, a behavior that also allows air intake to the tracheal system through the neighboring posterior spiracles (Hayashi and Kondo, 2018; Wu et al., 2003) (SupplFig. 4 B), and when larvae eventually exit the food source to pupariate (Wu et al., 2003) (SupplFig. 4 B). With their exposed nature, these sensory neurons may also integrate other inputs such as chemical cues or even light from the environment (Stewart et al., 2015; Vermehren-Schmaedick et al., 2011; Xiang et al., 2010).

Neuronal regulation of the hematopoietic system and other organs is an important paradigm in biology, which is starting to come to light in a variety of species (Kumar and Brockes, 2012). In *Drosophila*, neuronal regulation of hematopoiesis is well established. Embryonic-lineage plasmatocytes depend on sensory neurons for their survival, proliferation and localization (Gold and Brückner, 2014, 2015; Makhijani et al., 2017; Makhijani et al., 2011; Makhijani and Brückner, 2012). Activin-β, a TGF-β family ligand, is a key signal produced by active sensory neurons that promotes plasmatocyte proliferation and adhesion (Makhijani et al., 2017). Identification of the signal/s from active sensory cone neurons that trigger transdifferentiation will be the focus of intense future study. We postulate secreted factor/s, which could potentially, albeit at reduced efficiency, also act at a distance, promoting transdifferentiation of smaller numbers of crystal cells in the dorsal vessel associated hemocyte clusters (Leitao and Sucena, 2015) and segmental hematopoietic pockets. In this context it is interesting to note that in long term memory formation, Activin expression is induced downstream of calmodulin/CaMK/CREB signaling in both *Drosophila* and vertebrates (Inokuchi et al., 1996; Miyashita et al., 2012), suggesting potential parallels of this signaling cassette in neuron-induced blood cell transdifferentiation.

In vertebrates, functional links of hematopoiesis with sensory neurons or other sensing systems remain largely unknown, despite some aspects of bone marrow hematopoiesis and inflammatory responses being regulated by the autonomic nervous system (Hanoun et al., 2015; Pavlov and Tracey, 2012). Oxygen sensing could be an important regulatory factor in recently identified hematopoietic sites such as the vertebrate lung, which provides a microenvironment for limited blood cell progenitors and megakaryocytes that are active in platelet production (Lefrancais et al., 2017; Martin et al., 1983). The lung, like many other organs, also harbors tissue macrophages that proliferate in local microenvironments and bear evolutionary parallels with *Drosophila* embryonic-lineage plasmatocytes (Gold and Brückner, 2014, 2015; Perdiguero and Geissmann, 2016). Interestingly, the lung and airways are rich in vagal afferent nerves that sense chemical and mechanical cues (Chang et al., 2015; Mazzone and Undem, 2016), and neuroendocrine cells of the lung that sense oxygen early in life and later provide a microenvironment for airway epithelial cells (Caravagna and Seaborn, 2016; Cutz et al., 2007). Investigation of sensory neurons and other sensors may therefore open a new chapter in the regulation of vertebrate hematopoiesis, transdifferentiation, and immune cell fate and function.

## Materials and Methods

### Drosophila Strains

*Drosophila* drivers, reporters and related lines used were *HmlΔ-GAL4, UAS-GFP* (Sinenko and Mathey-Prevot, 2004); combination driver *HmlΔGAL4, UAS-GFP; He-GAL4* (Yang et al., 2015) (gift from Dan Hultmark), *lz-GAL4; UAS-GFP* (J. Pollock, Bloomington), *21-7-GAL4* (Makhijani et al., 2011; Song et al., 2007) *Gyc89Db-GAL4 (Morton et al., 2008)* (gift from David Morton); *BcF6-GFP* (Tokusumi et al., 2009) (gift from Robert Schulz); *BcF6-mCherry* (Tokusumi et al., 2009), *BcF2-GFP* (Tokusumi et al., 2009); *HmlΔ-DsRed* (Makhijani et al., 2011); and *tubGAL80ts* (McGuire et al., 2003). UAS lines used were *UAS-CD8-GFP* (Song et al., 2007); *UAS-Kir2.1* (Baines et al., 2001) (Bloomington); *UAS-TrpA1* (Bloomington); *UAS-Hid ala5* (Bergmann et al., 2002) (gift from Andreas Bergmann); *UAS-Gyc88E RNAi* (Bloomington). Control lines used were *w1118* (Bloomington) or *yw* (Bloomington). Unless otherwise stated, fly crosses were set and maintained at 25° Celsius. Crosses with the driver *Gyc89Db-GAL4* were performed at 29°C to enhance expression.

### Hemocyte Quantification

Total hemocyte quantification was performed essentially as described in (Corcoran and Brückner, 2020; Petraki et al., 2015). All fluorescently-marked hemocytes of single larvae were released into wells marked by a hydrophobic PAP pen (Beckman Coulter) on glass slides filled with 20-30 μL PBS. Cells were allowed to settle for 15-20 min, and were imaged by fluorescence tile scan microscopy on a Leica DMI4000B microscope with Leica DFC350FX camera and 20x objective. Cell numbers in images were analyzed by particle quantification using Fiji/ImageJ (Corcoran and Brückner, 2020; Petraki et al., 2015; Schindelin et al., 2012).

Crystal cell quantification was performed using fluorescent protein reporters or melanization. To quantify crystal cells in live animals, larvae were placed on a slide in a small drop of 10-15 µl PBS with a coverslip on top. Fluorescent crystal cells in each segment were manually counted, rolling the larva by gently moving the coverslip. For phagocytosis lineage tracing, fluorescent crystal cells were quantified ex vivo (see below). To quantify crystal cells based on their ability to melanize (due to their hallmark expression of functional Prophenoloxidase 1 and 2) larvae were placed in 250 µl of PBS in Eppendorf tubes and heated at 65° Celsius for 22 minutes in a heat block, and melanized (black) cells were quantified under a stereoscope by manual counting (Corcoran and Brückner, 2020; Rizki and Rizki, 1959).

Larvae were analyzed at various developmental times after egg laying (AEL) corresponding to the following size ranges: 1^st^ instar: ∼0.5-1.4mm (22-46h AEL; 2^nd^ instar: ∼1.5-2.6mm (47-77h AEL); 3^rd^ instar: ∼2.7- >3.5mm (from 78h AEL). For the crosses used, no developmental delays were observed; therefore selection of specified larval size ranges from 24h embryo collections was used in lieu of more tightly timed embryo collections.

### Manipulation of Neuronal Activity

To mimic sensory neuron stimulation, larvae were exposed to 10mg/ml carbamylcholine (carbachol, Sigma Aldrich) in fly food for 4 hours allowing direct cuticle contact with carbachol (Makhijani et al., 2017).

To activate specific neuron populations, the heat-inducible cation channel TrpA1 was ectopically expressed and transiently heat induced to mimic neuron activation (Hamada et al., 2008). Sopecifically, *TrpA1* crosses were set at RT and shifted to 29°C for 4h. Larvae were analyzed within 30 minutes following this period.

Sensory neuron silencing was achieved by transiently expressing a transgenic of the inward rectifying potassium channel Kir2.1 (Baines et al., 2001), under control of a sensory neuron specific GAL4 driver and a temperature-sensitive GAL4 inhibitor GAL80ts (McGuire et al., 2003); genotypes for neuronal silencing experiments were *21-7-GAL4, UAS-GFP, HmlΔDsRed/UAS-Kir2.1; tubGAL80ts/*+ with controls *21-7-GAL4, UAS-GFP, HmlΔDsRed/*+; *tubGAL80ts/*+. F1 from crosses were raised at 18° Celsius. To temporarily silence sensory neurons, larvae were shifted from 18° to 29° Celsius for 22 hours to destabilize GAL80ts, allowing for expression of Kir2.1, which hyperpolarizes sensory neuron preventing firing. Larvae were analyzed within 1 hour post-silencing.

### Phagocytosis Lineage Tracing

For lineage tracing of crystal cells that derive from actively phagocytic plasmatocytes, larvae with genotype *HmlΔGAL4, UAS-GFP; BcF6-mCherry* were injected with 69 nl of blue fluorescent FluoSphere Carboxylate-modified 0.2 um beads (Invitrogen) diluted 1:100 in PBS using a Nanoject injector (Drummond Scientific). After injection, larvae were placed on food with yeast for 4 hours. Hemocytes of individual larvae were then released into a well marked on a glass slide by PAP pen and filled with 20-30 µl PBS; hemocytes were allowed to settle for 15-20 minutes in a wet chamber (Corcoran and Brückner, 2020; Petraki et al., 2015). Cells were imaged with a Leica DMI4000 microscope with tilescan function. Cell quantification was conducted manually from a representative central field of each image (containing ∼150-300 plasmatocytes). For genotypes that give rise to substantially reduced numbers of crystal cells, crystal cells of the whole tilescan area were analyzed. Transdifferentiation was quantified by comparing ratios of crystal cells positive for blue particles/ total crystal cells (mCherry positive) for various experimental and control conditions. The same quantification for plasmatocytes (GFP positive) with blue beads relative to all plasmatocytes was conducted as an internal control for injection efficiency and phagocytic fitness.

### Hypoxia experiments

Hypoxia experiments were conducted using a hypoxic chamber (Biospherix, Inc., Laconia, NY). Oxygen concentrations were set as indicated, supplementing reduced O_2_ with N_2_. Incubations were performed at room temperature (RT) for 6h. Crystal cell melanization assays were conducted with larvae of the genotype *w*^*1118*^, and phagocytic lineage tracing and total hemocyte experiments used *Hml*Δ*-GAL4, UAS-GFP; BcF6-mCherry* larvae. For all experiments, 15-25 larvae of the desired age/size and genotype were placed on inactivated yeast paste in cell strainer snap cap tubes, allowing for rapid gas exchange (Corning #352235). Six cell strainer tubes were held by a fitted rack that was inserted into a large cylindrical polystyrene *Drosophila* population cage with steel mesh on one side (custom construction). During transport from the hypoxic chamber to the bench, population cages were sealed in a plastic bag to maintain hypoxic conditions up to the point when larvae were analyzed. Control samples were placed in an identical setup but left at normoxic conditions at comparable temperature (RT) and time (6h). Following hypoxia/normoxia conditions, assays were conducted as described in the respective paragraphs.

### Microscopy

Released hemocytes were imaged on a Leica DMI4000B microscope as described above. Live imaging of larvae was done as described previously, immobilizing larvae on an ice-cooled metal block (Makhijani et al. *Development* 2011). Imaging was performed using a Leica M205FA fluorescent stereoscope with DFC300FX color digital camera and Leica LAS Montage module, combining z-stacks into single in-focus images.

### Statistical Analysis

For all experiments numbers of larvae per genotype and condition are indicated in the Figure Legends. For each genotype and condition the mean and standard deviation were determined and significance was tested by 2-way ANOVA (Prism). For hemocyte counts over the course of larval development, regression analysis was performed (Excel). For phagocytosis lineage tracing, the mean and standard deviation of percentages of blue bead positive cells were determined and assessed by 2-way ANOVA (Prism). P-value cutoffs for significance were as follows: * = p < 0.05, ** = p < 0.01, and *** = p < 0.001. Pools of both male and female larvae were analyzed.

## Authors’ contributions

KB conceived and supervised the study. SC, AM and KB planned the experiments. SC, AM, JA, YH, DO, TJ, KK carried out the experiments. SC, AM, JA, YH, DO, TJ, KK and KB analyzed the data. KB wrote the manuscript with input from all authors.

## Acknowledgements

We thank D. Morton, R. Schulz, U. Banerjee, A. Bergmann, C. Evans, D. Hultmark, Y.N. Jan, L. Kockel, the Bloomington Stock Center, and TRiP for fly stocks. We are grateful to Khalida Sabeur and Manideep Chavali for help to access and use hypoxia chamber equipment. Thanks to Ava Brückner-Kockel for advice on regression analysis. We thank members of the Brückner lab for discussion and feedback on the manuscript.

## Supplemental Information

**Supplemental Figure 1.**
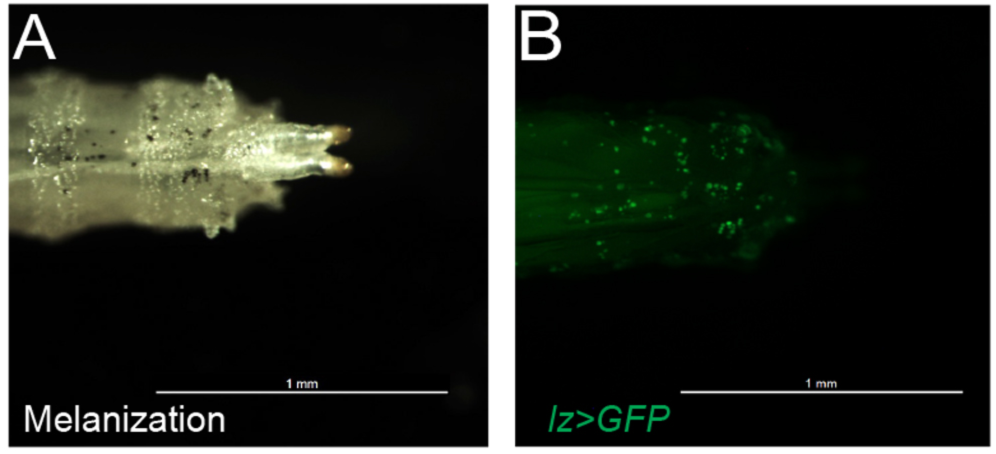
Methods of crystal cell labeling. (A) Heat induced melanization of crystal cells. Genotype is *lz-GAL4;UAS-GFP* (B) Fluorescent reporter labeling to visualize, and quantify, crystal cells. Genotype is *lz-GAL4;UAS-GFP*. Note that the labeled crystal cell pattern by both methods is very similar.

**Supplemental Figure 2.**
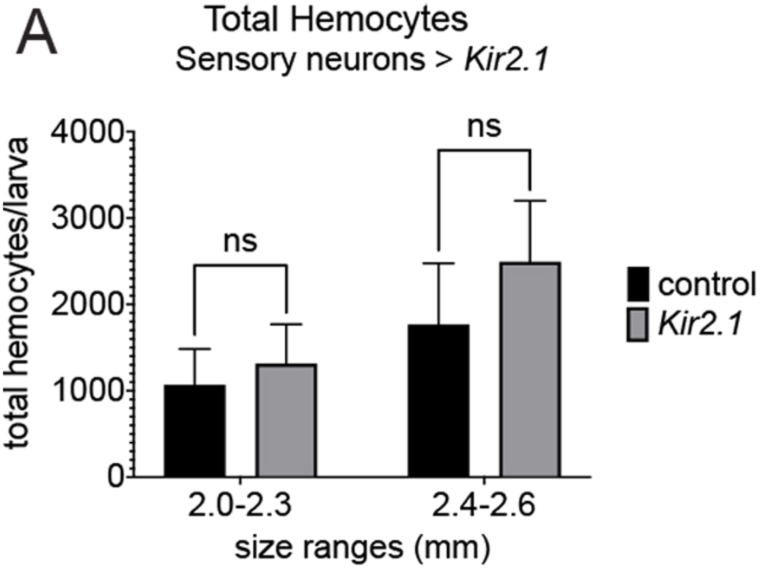
Limited transient sensory neuron silencing does not affect total hemocyte numbers. (A) Transient silencing of sensory neurons, quantification of total hemocytes; genotypes are *21-7-GAL4, UAS-CD8-GFP, Hml*Δ*-DsRed/ UAS-Kir2.1; tubGAL80ts/* +, n=12 and control *21-7-GAL4, UAS-CD8-GFP, Hml*Δ*-DsRed/* +; *tubGAL80ts/* +, n=13. Larvae induced at 29°C for 22h. Mean and standard deviation, two-way ANOVA

**Supplemental Figure 3.**
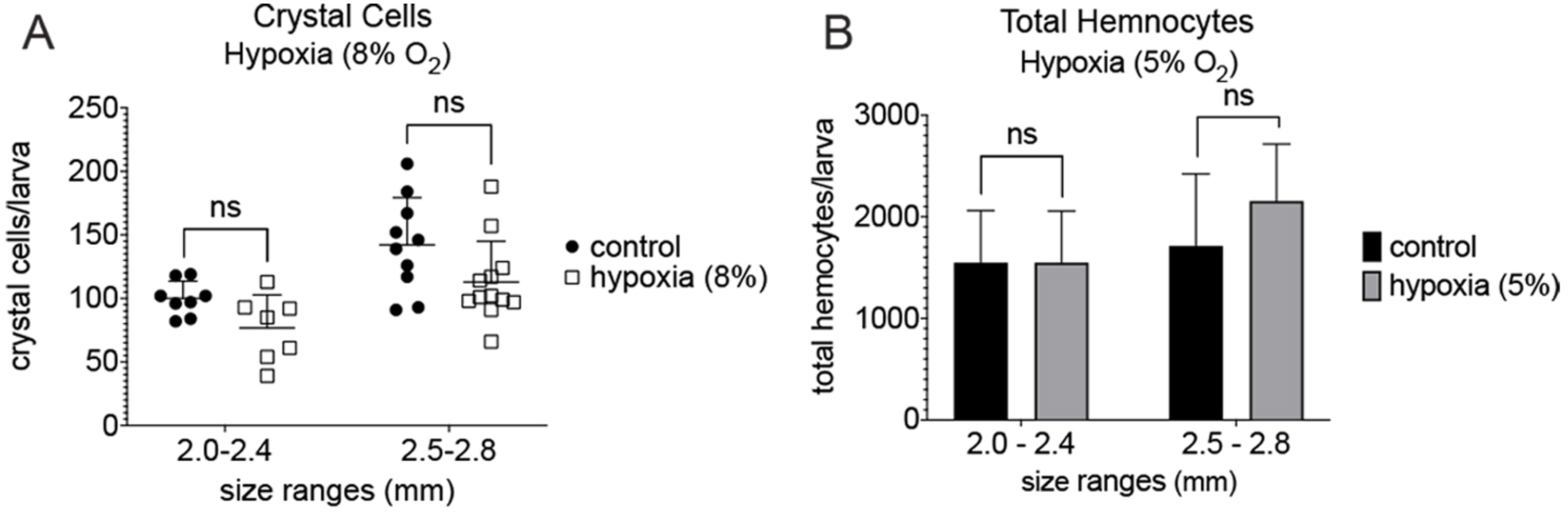
Hypoxia affects crystal cell counts but not total hemocyte numbers. (A) Effect of 8% hypoxia on crystal cell number determined by melanization; genotype is *w1118*; hypoxia n=19 and normoxia control n=18. Individual value plot with mean and standard deviation, two-way ANOVA. (B) Total hemocyte number under conditions of hypoxia (5% O2) and normoxia; genotype is *Hml*Δ*-GAL4, UAS-GFP; BcF6-mCherry*; hypoxia n=14 and normoxia control n=14. Mean and standard deviation, two-way ANOVA.

**Supplemental Figure 4.**
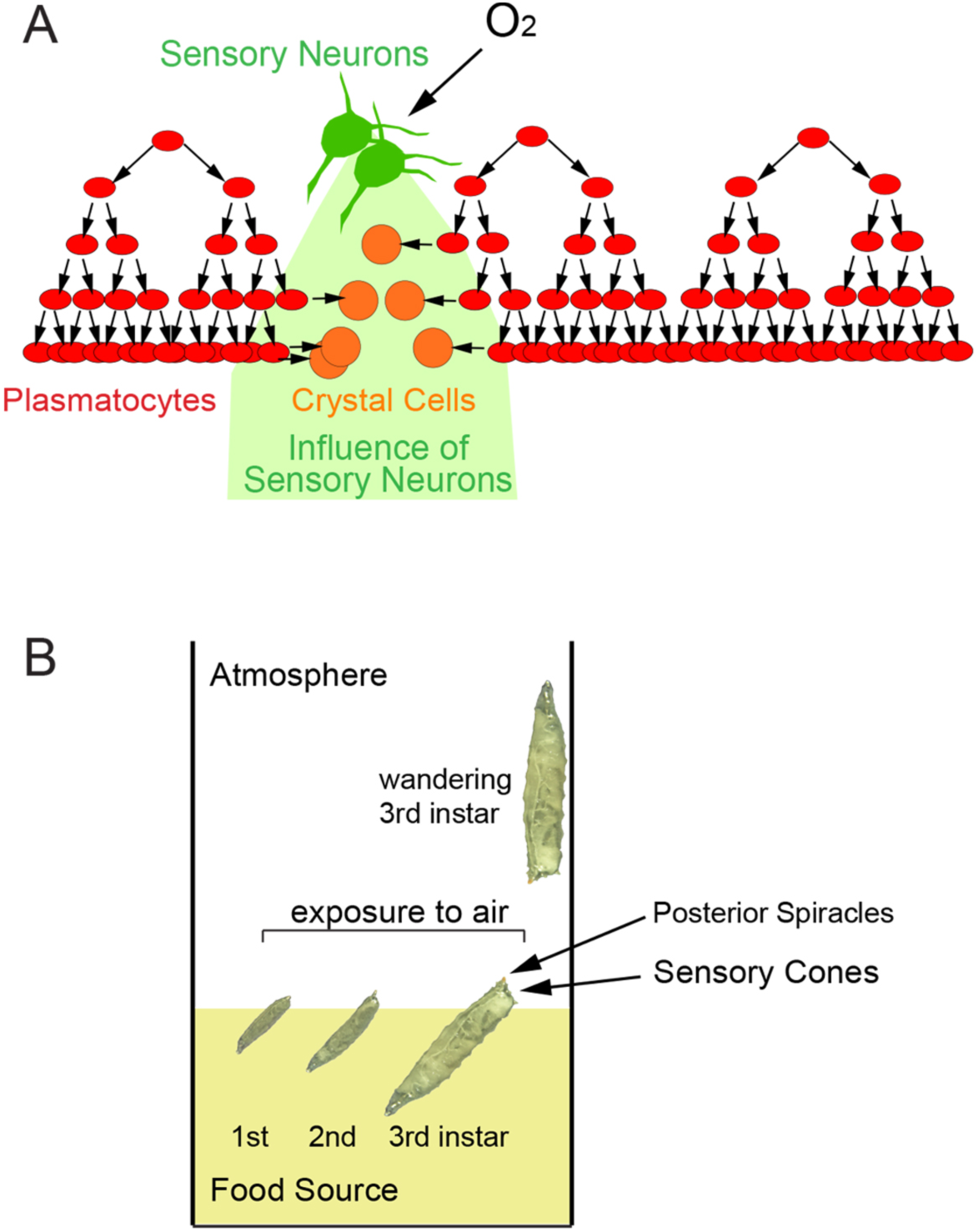
Models. (A) In the presence of oxygen, active sensory cone neurons produce signal/s that drive transdifferentiation of plasmatocytes to crystal cells. According to this model, transdifferentiation is triggered by a, likely secreted, sensory neuron signal, therefore plasmatocytes in anatomical proximity to the sensory cone neurons are most likely to convert to crystal cells. (B) Model illustrating exposure of the caudal end of *Drosophila* larvae including the sensory cones and posterior spiracles to the air, while burying in food. Mature larvae leave the food in preparation of pupariation, now fully exposed to the air.

